# Manipulating Oxalate Decarboxylase Provides the Basis of Antilithic Therapy by Acting on the Gut Microbiota

**DOI:** 10.1101/2022.10.14.512337

**Authors:** Fang Wu, Yuanyuan Cheng, Jianfu Zhou, Peisen Ye, Xuehua Liu, Lin Zhang, Rongwu Lin, Songtao Xiang, Zhongqiu Liu, Caiyan Wang

## Abstract

A high concentration of oxalate is associated with an increased risk of kidney calcium oxalate (CaOx) stones, and the degradation of exogenous oxalate mainly depends on oxalate-degrading enzymes from the intestinal microbiome. We found that Zinc Gluconate supplement to patients with CaOx kidney stones could significantly improve the abundance of oxalate metabolizing bacteria in human body through clinical experiments on the premise of simultaneous antibiotic treatment and the imbalance of *Lactobacillus* and OxDC was involved in CaOx kidney stones through clinical sample analysis. Then, we identified that Zn^2+^ could be used as an external factor to improve the activity of OxDC and protect *Lactobacillus*, achieved the preventive effect on rats with stones aggravated by antibiotics. Finally, by analyzing the three-dimensional structure of OxDC and some *in vitro* experiments, we propose a hypothesis Zn^2+^ increases the metabolism of oxalate in humans through its positive effects on *Lactobacillus* and OxDC to reduce CaOx kidney stone symptoms in rats.

**IMPORTANCE:** Urinary stone disease is one of the most common urological disorders, and 70%-80% of urinary stones are calcium oxalate (CaOx) stones. We found the structural basis and metabolic mechanism by which oxalate decarboxylase metabolizes oxalate were elucidated, and Zn^2+^ was illustrated to have therapeutic effects on CaOx stones by improving the tolerance of Lactobacillus to antibiotics. According to that, proper Zn^2+^ levels in the diet, the consumption of more probiotic food and avoidance of the antibiotic overuse might be desirable measures for the prevention and treatment of kidney stones.

**Graphical Abstract:** 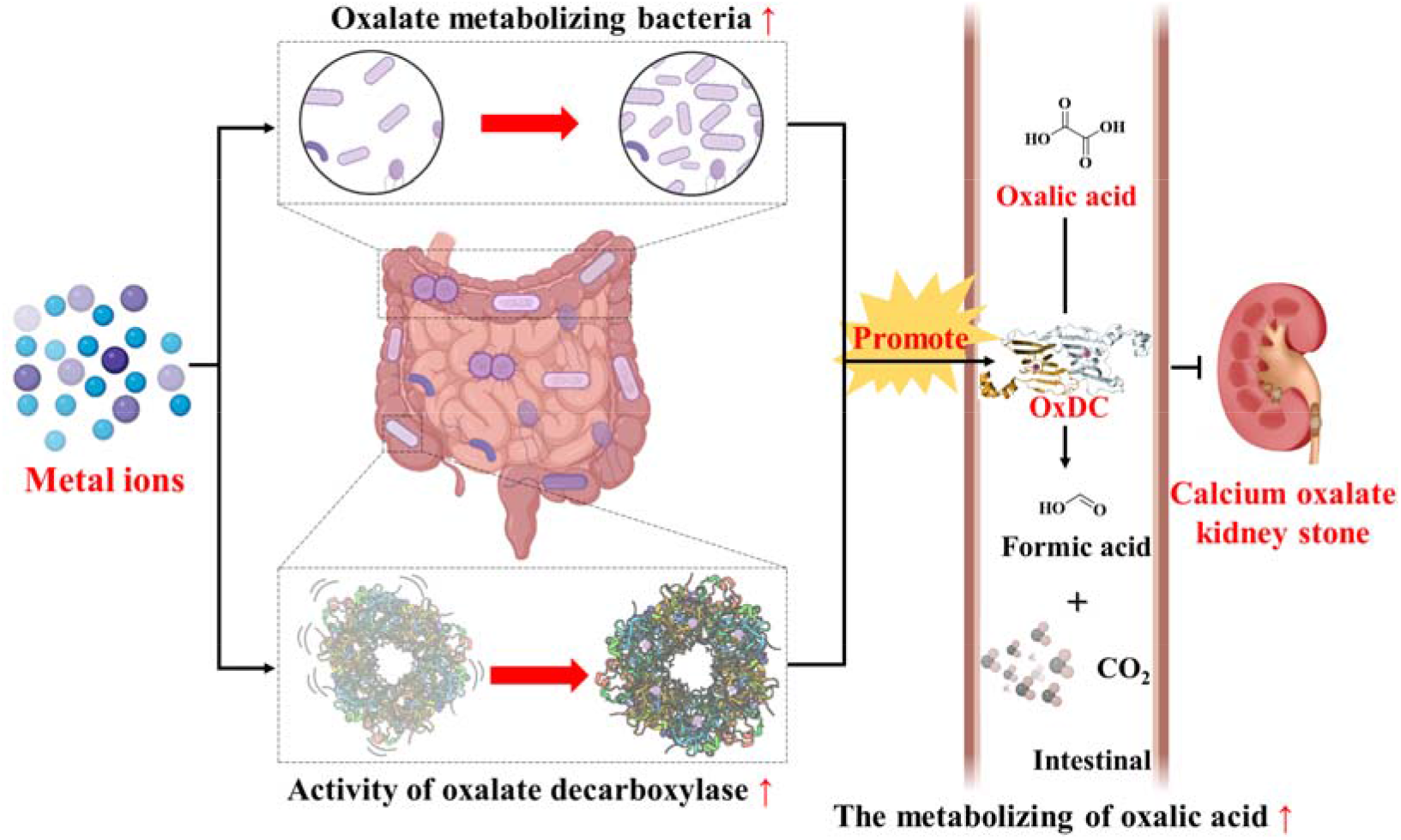

## Introduction

Urinary stone disease is one of the most common urological disorders (1), and 70%-80% of urinary stones are calcium oxalate (CaOx) stones (2-5). Clinically, patients with CaOx stones often exhibit hyperoxaluria and hypercalciuria (6); when oxalate excretion exceeds 40-45 mg per 24 h, the patient will be diagnosed with high oxaluria and is at risk for CaOx stones (7). The curved distal renal tubules exposed to a high concentration of oxalate over a long time are liable to form small crystalline nuclei that damage kidney epithelial cells and promote the formation of initial plaques. Eventually, plaques detach from the lumen, form a large CaOx stone and cause inflammation in the kidney (8, 9). Thus, the high content of oxalate in the human body is a major causal driver to kidney stones.

The human body does not contain any enzyme system for oxalate metabolism (10). Approximately 80% of exogenous oxalate is metabolized depending on oxalate-degrading bacteria (11, 12). Thus, the oxalate level in the kidneys is mainly modulated by the intestinal flora responsible for metabolizing oxalate. Moreover, long-term use or abuse of antibiotics causes intestinal flora disorder. In particular, the reduction of oxalate-metabolizing bacteria would significantly increase the risk of forming CaOx stones (13, 14). Accordingly, maintaining the balance of the intestinal flora in the human body and increasing the number of oxalate-degrading bacteria positively effect on patients with CaOx stones.

At present, although the treatment of kidney stones has evolved to advanced minimally invasive surgery, the recurrence rate of stones is still in a high level (15, 16). Therefore, it is important to discover new effective treatment strategies for CaOx stones patients. *Lactobacillus* strains are commonly used in dairy products and medical probiotic formulations due to their efficiency and safety (17-20). Recently, Campieri *et al* (21-25) have found that the oxalate content and stone size were significantly reduced in patients with hyperoxaluria after the *Lactobacillus’s* administration. The degradation of oxalate by *Lactobacillus* is attributed to the expression of oxalate decarboxylase (OxDC) (26). The therapeutic effect of OxDC was further confirmed by the CaOx stones rat models (27). Besides, *Lactobacillus* can strengthen the intestinal barrier by regulating the expression of tight junction protein and the body’s immune response (28, 29), playing an important role in maintaining the balance of intestinal flora (30). Given the advantages of *Lactobacillus* and OxDC mentioned above, it has become a promising and potential strategy for the treatment of CaOx stones patients. Nevertheless, the oxalate’s metabolic mechanism by OxDC is still largely unknown, limiting the widespread use of *Lactobacillus* and OxDC in the clinic.

Zinc is the second most abundant essential trace element that plays a vital role in maintaining the normal function of the human body (31). Zinc serves as a metal cofactor for more than 300 enzymes (32), and has an anti-inflammatory function (33). As probiotics, *Lactobacillus* has a good zinc enrichment effect, which increases their antibacterial activity in the intestine (34). If zinc can improve the ability of *Lactobacillus* to degrade oxalate in the intestine or enhance its viability, it would play a positive role with the treatment of CaOx stones.

In this study, we learned that in clinical treatment experience supplementation of Zinc Gluconate for CaOx kidney stone patients would improve the gut microflora diversity and the abundance of of *Lactobacillus*. Thus, we explored the relationship among the Zn^2+^, *Lactobacillus* and oxalate metabolism. Firstly,The protective effect of Zn^2+^ on *Lactobacillus* was discovered *in vitro*, and it was further confirmed that Zn^2+^ had a relieving effect on CaOx kidney stone symptoms aggravated by antibiotics *in vivo*. Secondly, it was found that the content of OxDC in patients with CaOx kidney stones was abnormal significantly, suggesting that there may be abnormal oxalate metabolism in patients, and then based on the co-crystal structure of OxDC and formic acid, combined with enzyme activity and cell assays, we elucidated the mechanism of oxalate degradation by OxDC. Finally, we found that Zn^2+^ could improve the stability and oxalate-degradation activity of OxDC and determined its specificity for OxDC and *Lactobacillus*. In a word, we demonstrated that Zn^2+^ supplementation may alleviate symptoms in patients with CaOx kidney stones by increasing the enzymatic activity of OXDC as well as the abundance of *Lactobacillus*.

## Results

### Supplementation of Zinc Gluconate on gut microflora for patients with CaOx kidney stones

The excessive intake of oxalate and its metabolic disorders are key triggers of CaOx kidney stones formation. The oxalate metabolism is mainly dependent on four types of intestinal bacteria in human, including *Oxalobacter, Lactobacillus, Enterobacteriaceae*, and *Bifidobacterium*. In clinical treatment, we found feces in patients with CaOx kidney stones was improved after Zinc Gluconate therapeutics. We recruited 40 subjects and divided them into four cohorts, i.e.: non-kidney stones group (n=10), CaOx kidney stones group (n=10), CaOx kidney stones treated with antibiotic group (n=10), and CaOx kidney stones treated with antibiotic and Zinc Gluconate group (n=10). The feces of subjects were used as samples to determine the abundance of several important intestinal flora in human intestinal (**Figure 1A**). Analysis of intestinal microbes showed that the abundance of *Bacteroidota* and *Actinobacteriota* changed significantly. However, the change of human beneficial bacteria *Firmicutes* was not obvious in four groups (**Figure 1B**). Therefore, we further investigated *Lactobacillus* in *Firmicutes*, it was found that the abundance of *Lactobacillus* decreased significantly in the CaOx kidney stones group but increased significantly after Zinc Gluconate supplementation. At the same time, we also measured the abundance of two other oxalate metabolizing bacteria in human intestine, which was similar to the trend of *Lactobacillus* (**Figure 1C**). These results showed that administration of Zinc Gluconate supplement to patients with CaOx kidney stones could significantly improve the abundance of oxalate metabolizing bacteria in human body, especially with the premise of simultaneous antibiotic treatment. Similarly, we also detected changes in the diversity of intestinal flora, and the types of intestinal flora in patients with kidney stones were significantly reduced. After antibiotic-based treatment, the species of bacteria continued to decline, but after the adjuvant treatment with Zinc Gluconate, the number of species of bacteria almost returned to a normal level (**Figure 1D**).

**Figure 1.**
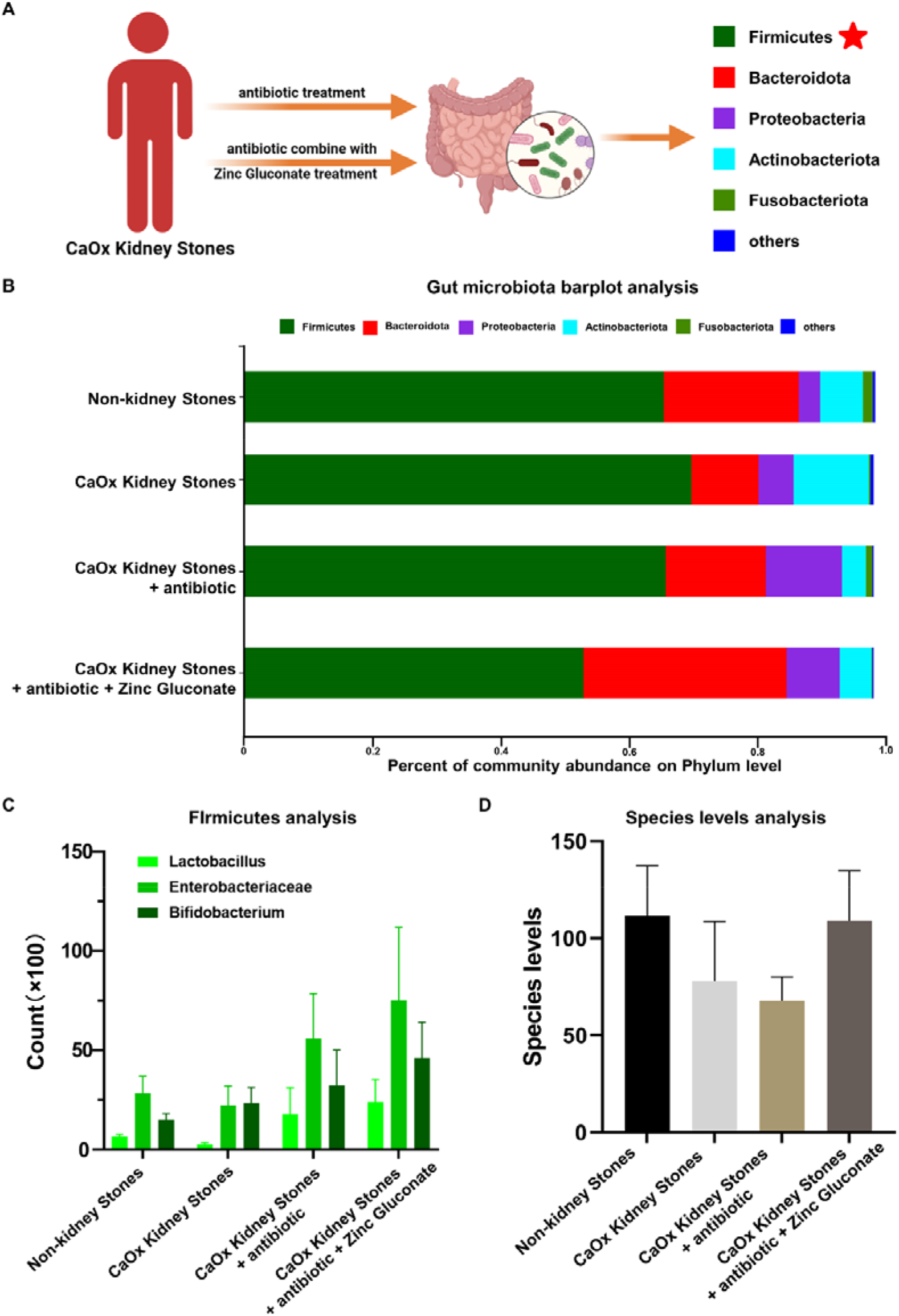
Clinical data analysis of the effects of Zinc Gluconate on the patients with CaOx stones. (A) Four cohorts were devised in this study: the antibiotic treatment, the antibiotic combines with Zinc Gluconate, the patients with CaOx Kidney stones and the patients with non-kidney stones. The last two groups were not shown in the figure. (B) Bar plot analysis of several major intestinal microbiota in the human intestinal. (C) The analysis the abundance of several *Firmicutes* in four groups. (D) The analysis the species levels of bacterial composition in four groups.

### The protection of Zn^2+^ on *Lactobacillus* and the therapeutic effect on rats with CaOx stones

According to clinical investigation, the abuse of antibiotics can lead to an imbalance of the human intestinal flora, so it is often regarded as a potential kidney stones trigger (35). *L. casei* is a kind of probiotics belonging to the genus *Lactobacillus* commonly used in life. Therefore, to screen out which antibiotic is extremely sensitive to *L. casei*, we used seven common antibiotics to interfere with the survival of *L. casei* (**Figure 2A**) and found that *L. casei* had the strongest sensitivity to ampicillin (Amp), chloramphenicol, and penicillin. Finally, we chose Amp as an inhibitor of *Lactobacillus* in subsequent assays. Then we tested the effect of different metal ions on intestinal flora. As expected, we found that Zn^2+^ could promote the growth of flora (**Figure 2B**). With the treatment with Amp, Zn^2+^ addition was significantly increased the survival rate of *L. casei* compared with other common divalent metal ions (**Figure 2C**). Since excessive oxalic acid inhibits the growth of *Lactobacillus*, we subsequently investigated the effect of Zn^2+^ on the tolerance of *Lactobacillus* and *L. casei* towards oxalic acid by adding Zn^2+^ to DeMan Rogosa-Sharpe (MRS) medium with the supplement of oxalic acid. The impact of Zn^2+^ on the survival of *Lactobacillus* and *L. casei* under the Amp condition was also investigated similarly. The addition of Zn^2+^ increased the tolerance of *L. casei* to oxalic acid (**Figure 2D**). The assay results also showed that Zn^2+^ could hold back the inhibition effect of oxalic acid or Amp on *L. casei*. The addition of Zn^2+^ could promote proliferation and increase survival rate of *L. casei*. According to literature reports, Zn^2+^ reusation of increased *E*.*coli* by relieving oxidative stress, and further promote the transcription of AdcA and AdCB by regulating the transcription inhibitor AdcR, thereby promoting the gene transcription translation of Streptococcus and increasing bacterial proliferation (36,37). These results indicated that Zn^2+^ may achieve protective effect by regulating the oxidative stress and transcriptional translation process of *L. casei* (**Figure 2E**).

**Figure 2.**
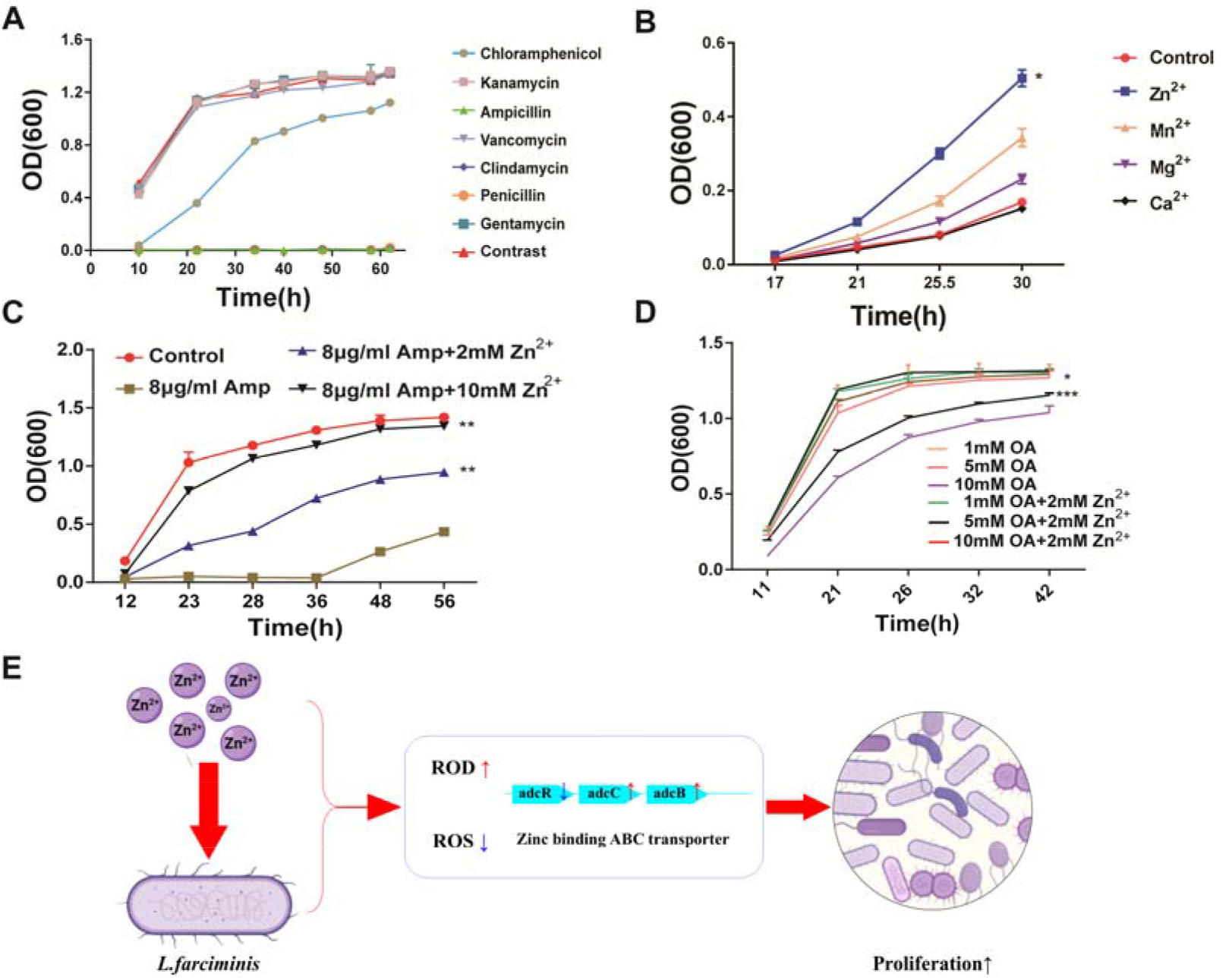
The protection of Zn^2+^ on *Lactobacillus* and the therapeutic effect on rats with CaOx stones. (A) The influence of seven common antibiotics on the growth of *L. casei*. (B) The influence of four common divalent metal ions (Zn^2+^, Mn^2+^, Mg^2+^, Ca^2+^) on the ability of lactobacilli to tolerate Amp. (C) The survival of *L. casei* under antibiotic stress after the addition of Zn^2+^. (D) Effect of Zn^2+^ on the tolerance of *L. casei* to oxalic acid. (E) Schematic diagram of the mechanism by which Zn^2+^ protects *Lactobacillus* and promotes its proliferation. All assays were replicated more than three times. * indicates *P*<0.05; ** indicates *P*<0.01; *** indicates *P*<0.001.

### The effect of Zn^2+^ on the inflammatory response and the degree of kidney damage

We further intent to explore whether Zn^2+^ had a certain effect on CaOx stones *in vivo*. Based on the H&E staining results, the rats with stones aggravated by antibiotics had a larger CaOx stone area compared to the control group. However, after the addition of Zn^2+^, the stone area reduced significantly compared to antibiotics aggravated stones rats (**Figure 3A**). To better evaluate the degree of damage of kidney stones in rats, we used osteopontin (OPN) as an indicator. OPN is one of the main organic components of CaOx stone matrix. The abnormal high content of OPN could accelerate the release of adhesion factors, increase kidney damage, and promote kidney inflammation (36)(**Figure 3B**). To further verify the therapeutic effect of Zn^2+^ on rat models with stones exacerbated by antibiotics, we measured inflammation-related indicators. We found that OPN and CD44 mRNA levels in the kidney of the stone group and the “Amp+Stone” group were significantly increased compared with the control group (**Figure 3C**). Secondly, urea nitrogen in serum increased significantly, while urea nitrogen excreted in the urine reduced remarkably in the “Amp+Stone” group, which indicated that the kidney was damaged, but given zinc supplements group did not change significantly (**Figure 3D**). Finally, we also tested the expression level of inflammatory factors (TNFα, IL-1β) and zinc supplements administration suppressed the mRNA level of TNFα and IL-1β (**Figure 3E**). These results further illustrated that Zn^2+^ had a certain therapeutic effect on rats with stones exacerbated by antibiotics.

**Figure 3.**
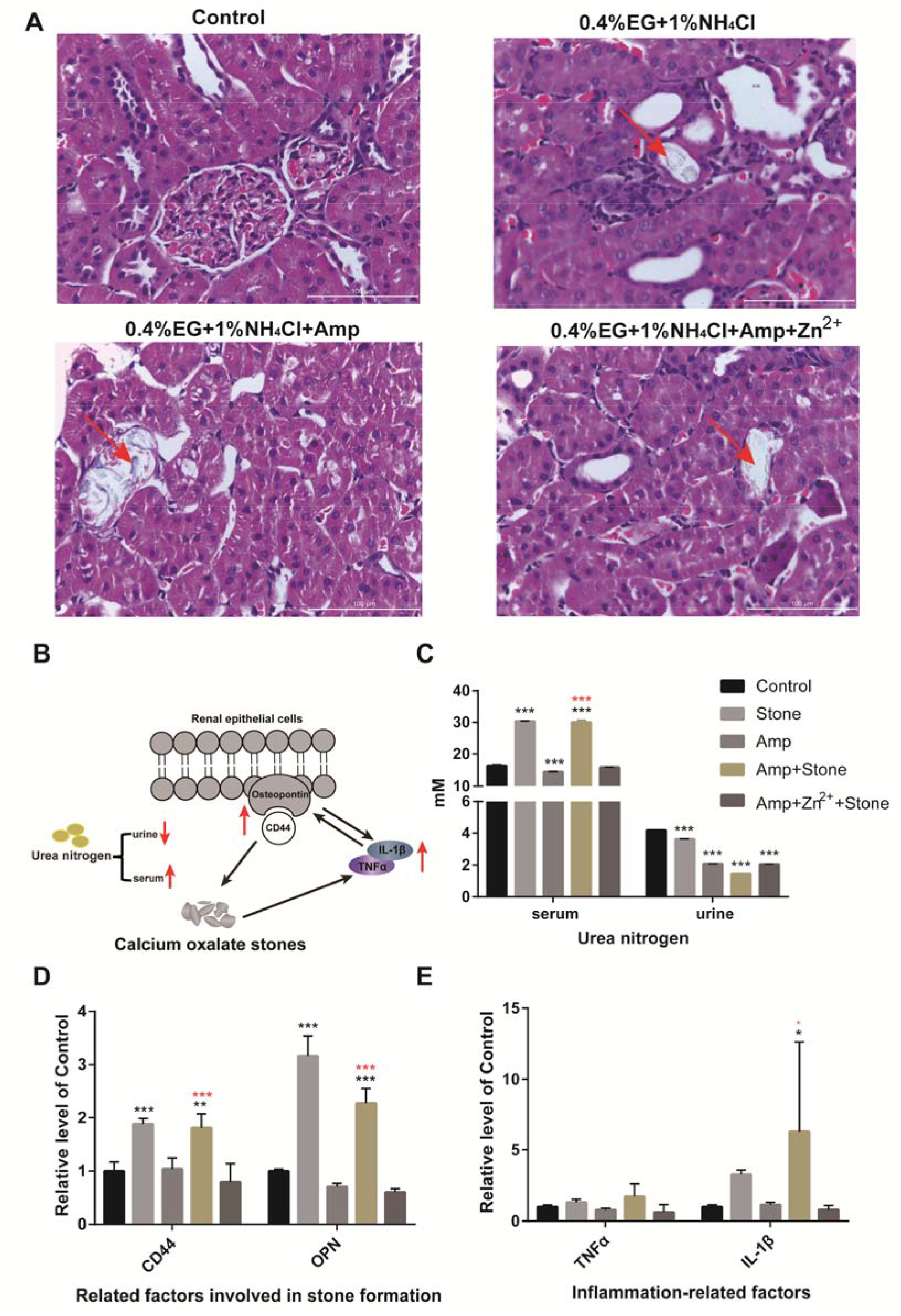
The effect of Zn^2+^ on the inflammatory response and the degree of kidney damage in rats with CaOx stones. (A)Results of hematoxylin-eosin staining of rat kidney tissue after 11 d of modeling. The control group was given the same volume of 0.9% saline. The red arrow shows the sites of stone growth. (B) Diagram of the relationship between inflammatory factors, osteopontin and CaOx stones. (C) Changes of urea nitrogen in serum and urine in rats with CaOx stones. (D) Changes in mRNA levels of osteopontin and its receptor (CD44) in kidney tissue. (E) The expression level of inflammatory factors (TNFα,IL-1β) in the kidney tissue of rats with CaOx stones were detected by qRT-PCR. The black * means compared with the control group, and the red * means with Amp+Stone as the reference group. * indicates *P*<0.05; ** indicates *P*<0.01; *** indicates *P*<0.001.

### Imbalance of OxDC in CaOx kidney stones patients and the overall structure of *L. farciminis* OxDC

Previously, we demonstrated that Zn^2+^ could promote the abundance of beneficial intestinal microflora and CaOx metabolism. Since *Lactobacillus* was reported to express OxDC to degrade oxalate (26), which had great potential for the treatment of CaOx stones, we further evaluated the mRNA level of OxDC in two groups. Compared with that in the non-kidney stones patients’ group, the mRNA level of OxDC reduced sharply in the CaOx kidney stones patients’ group (**Figure 4A**), consistent with the low level of *Lactobacillus* in the CaOx kidney stones patients. These results implied that both OxDC and *Lactobacillus*’s concentration might be closely related to the formation of CaOx stones.

**Figure 4.**
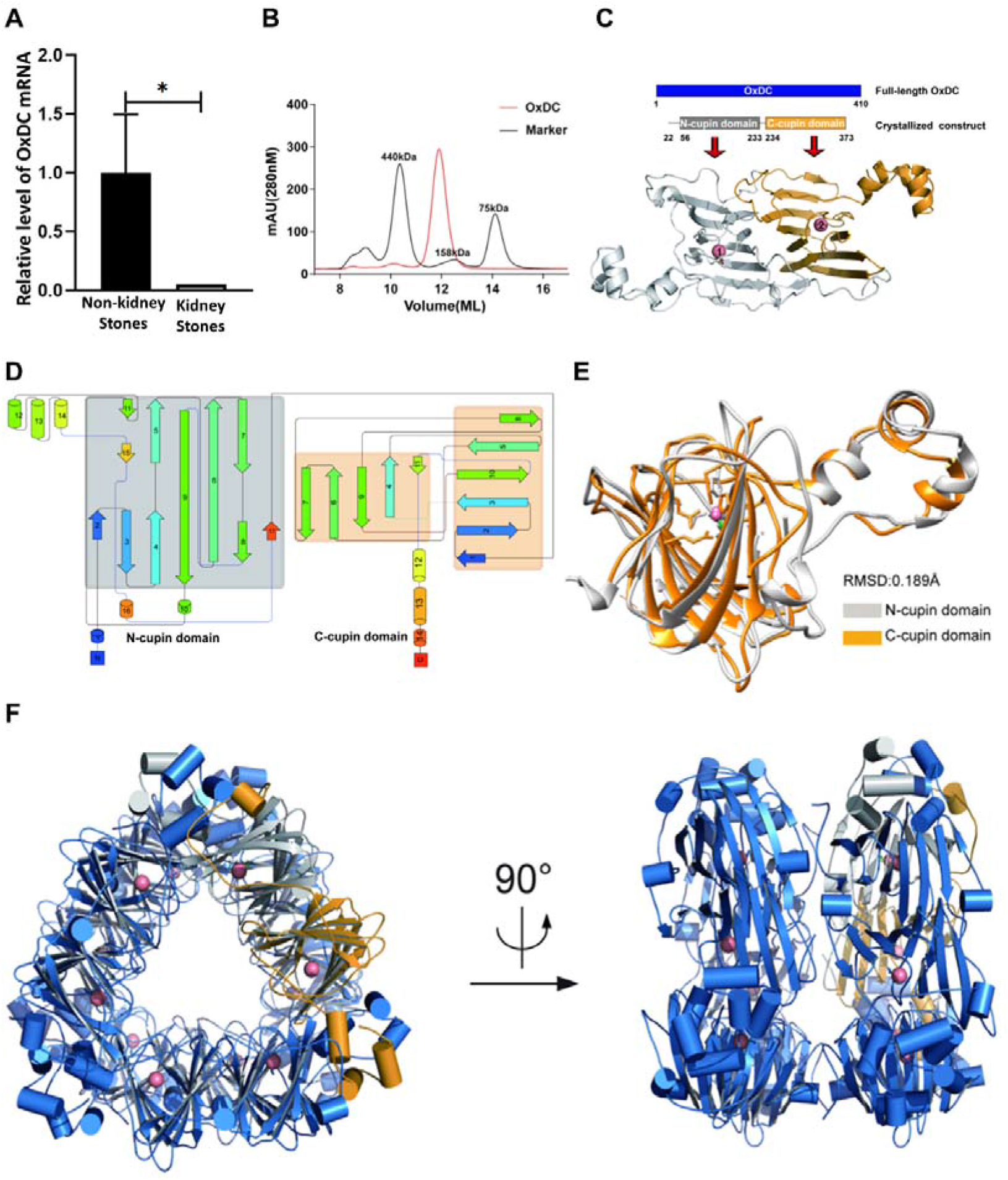
Association of oxalate-degrading bacteria and OxDC with CaOx stone formation in kidney stones patients and the crystal structure of *L. farciminis* OxDC. (A) The mRNA level of OxDC in the non-kidney stones and CaOx kidney stones patients. The number of patients in the non-kidney stones group and CaOx kidney stones group is 25 and 27, respectively. Measurement data were represented as the mean ± SEM. Unpaired t-test was conducted to assess data with a normal distribution and equal variance. (B) Chromatogram from size-exclusion chromatography analysis. The black line represents the chromatogram of three standard molecular weight proteins (440, 158, and 75 kDa), and the OxDC chromatogram is indicated with the red line. (C) Domain distribution and structure of a single subunit of OxDC. The N-terminal and C-terminal cupin domains are colored gray and orange, respectively. Mn^2+^ is presented as pink spheres. (D) OxDC’s topology diagram. The shaded color indication is consistent with Figure A. The columnar is represented as α-helix, and the arrow stands for β-sheet. (E) The structural similarity of the two functional domains. The N-terminal and C-terminal cupin domains are colored gray and orange, respectively. The software Pymol was used for structural Alignment. (F) The hexamer structure of OxDC in cartoon mode (front view and view after rotating 90° along the Y-axis). * indicates *P*<0.05; ** indicates *P*<0.01; *** indicates *P*<0.001.

Considering the importance of *L. farciminis* OxDC in the degradation of CaOx kidney stones, we intended to explore the metabolic mechanism of OxDC started from protein structure. We constructed the expression system of OxDC *in vitro*, and the chromatographic peak of OxDC showed a single symmetrical peak at the elution position between markers 440 kDa and 158 kDa (**Figure 4B**). The OxDC gene of *L. farciminis* encodes 410 amino acids (aa), which contains a N-terminal cupin domain (aa 56-233) and C-terminal cupin domain (aa 234-373). Fortunately, our crystal structure captures the active state of OxDC, which contains one metabolite formic acid in the N-terminal cupin domain (**Figure 4C left**). However, no electron density for formic acid was observed in the C-terminal cupin domain (**Figure 4C right**), The topological structure of the two domains is similar, with each consists of 11 β-sheets (**Figure 4D**). The RMSD value is 0.189 Å between the cupin domains, superimposition of both domains revealed no noticeable backbone movement or meaningful shift of the ligand position (**Figure 4E**). This phenomenon was highly likely due to the method of preparation of the crystal of the enzyme-substrate complex. The crystal structure of OxDC in complex with formic acid was resolved to a resolution of 2.79 Å by molecular replacement (**Table 1**). The structure of OxDC belongs to space group *P*_*21*_, with each asymmetric unit containing six OxDC molecules. This assembly state in crystal structure agreed with the chromatographic peak of OxDC. The overall structure of OxDC retains the conserved fold and sandwich morphology of the cupin family. In terms of internal interactions, the N-terminal of the upper layer of monomers were attached to the C-terminal of the lower layer and in the same layer, each pair of adjacent subunits were also joined by an interaction between the N-terminal and C-terminal domain, forming a stable triangular-like spatial conformation. A hollow cylindrical intermediate structure could be observed by structure rotating 90° along the Y-axis (**Figure 4F**). Metal ion was dispensable for cupin family proteins to carry out diverse biological functions (37), in OxDC structure, there were twelve endogenous Mn^2+^ evenly distributed in the two domains of each subunit, which were crucial for metabolizing the oxalate.

**Table 1.** Data collection and refinement statistics.

| Data collection |  |
| --- | --- |
| Space group | P21 |
| a, b, c (Å) | 92.96, 122.84, 115.22 |
| $\alpha, \beta, \gamma$ (°) | 90.00, 92.47, 90.00 |
| Resolution (Å) | 48.00-2.79 (2.84-2.79)<sup>a</sup> |
| CC<sub>1/2</sub> | 0.944 (0.810) |
| R<sub>merge</sub><sup>b</sup> | 0.141 (0.751) |
| I/$\sigma$(I) | 19.89 (1.94) |
| Completeness (%) | 96.94 (97.01) |
| Redundancy | 6.3 (6.0) |
| Refinement |  |
| Resolution (Å) | 48.00-2.79 (2.79-2.84) |
| No. reflections | 62185 (1795) |
| R<sub>work</sub><sup>c</sup>/R<sub>free</sub><sup>d</sup> | 0.197/0.229 |
| No. atoms |  |
| Protein | 16035 |
| Ligand | 18 (FMT),12 (MN) |
| Water | 248 |
| B-factors (Å<sup>2</sup>) |  |
| Protein | 35.8 |
| Ligand | 32.6 (FMT),36.5 (MN) |
| Water | 40.8 |
| R.m.s deviations |  |
| Bond lengths (Å) | 0.007 |
| Bond angles (°) | 1.109 |
| Ramachandran favored (%) | 97.37 |
| Allowed | 2.33 |
| Outliers (%) | 0.30 |

<sup>a</sup>Values in parentheses are for the highest-resolution shell. <sup>b</sup>R<sub>merge</sub> = $\Sigma |(I - \langle I \rangle)| / \Sigma I$ , where I is the observed intensity. <sup>c</sup>R<sub>work</sub> = $\Sigma_{hkl} ||F_o| - |F_c|| / \Sigma_{hkl} |F_o|$ , calculated from working data set. <sup>d</sup>R<sub>free</sub> is calculated from 5.0% of data randomly chosen and not included in refinement.

### Key residues for oxalate degradation by OxDC

OxDC degrades oxalic acid to formic acid and carbon dioxide. Based on analysis of conservation of the OxDC protein sequence and the structure of the protein in complex with formic acid, we found that each of the Mn^2+^ binding sites was coordinated by two highly conserved glutamate residues and three histidine residues in the cupin fold (**Figures 5A-C**). In the active conformation of OxDC, the formic acid molecule near Mn^2+^ interacted with the E162 in the N-terminal cupin domain, which was consistent with the previously reported state of *B. subtilis* OxDC (38).

**Figure 5.**
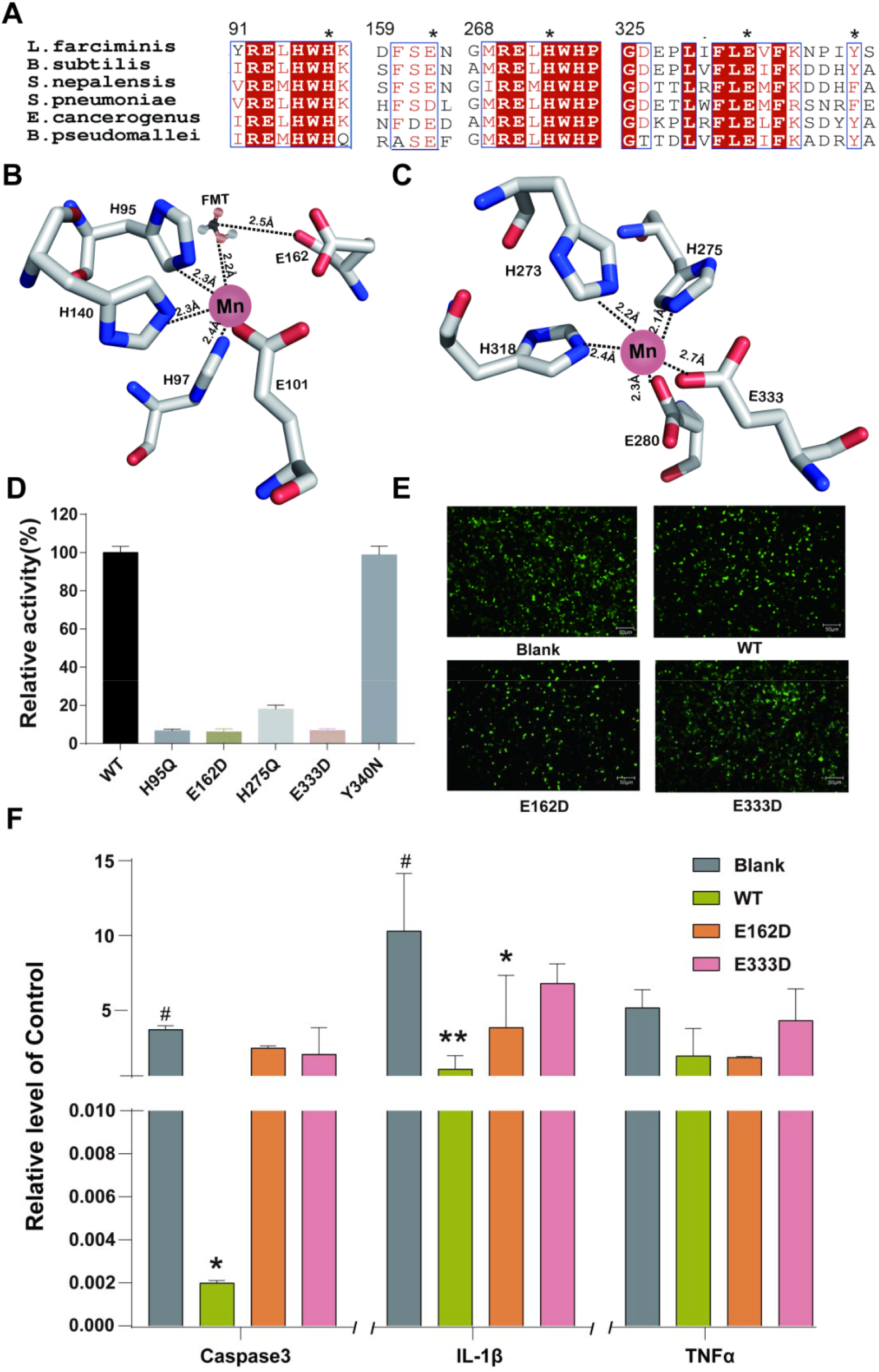
Discovery and verification of key residues in the active pocket. (A) Multiple amino acid sequence alignments between OxDC proteins from different species, including key fragments of two domains (red indicates highly conserved residues, asterisks indicate the five selected mutation sites). The GenBank identifiers (GI) for these species are 908212467 (*L. farciminis*) and 760455501 (*B. subtilis*), 1437302606 (*S. nepalensis*), 814506590 (*S. pneumonia*e), 1330860547 (*E. cancerogenus*), 126221798 (*B. pseudomallei*), respectively. (B) Schematic diagram showing coordination between key residues in N-cupin domains and Mn^2+^. FMT stands for formic acid. (C) Coordination between key residues in C-cupin domains and Mn^2+^. FMT is shown as a ball-and-stick model. (D) The enzyme activity of mutants and WT are verified *in vitro*. (E) Photographs taken under a fluorescence microscope after the transfection of exogenous plasmid (pEGFP, WT-pEGFP, E162D-pEGFP, E333D-pEGFP) into HEK293 cells. (F) Expression of apoptosis and inflammatory factors (Caspase3, IL-1β, and TNFα) in HEK293 cells. * indicates *P*<0.05; ** indicates *P*<0.01; *** indicates *P*<0.001.

We selected five conserved residues (H95, E162, H275, E333, and Y340) from the N and C-terminal cupin domains to investigate their role for the oxalate-degradation activity of OxDC by using oxalic acid as substrate with the methylene blue-potassium dichromate system. Compared with the wild type (WT) OxDC, the activity of mutants (H95Q, E162D, H275Q, and E333D) all severely harmed, confirming the crucial role of these four residues (**Figure 5D**). Mutant Y340N was equally active as WT OxDC. It was likely that the Y340 residue did not participate in the catalytic activity of oxalic acid, and it was not structural conservation among these five residues. Notably, according to previous literature reports (39), the N-cupin domain was a key site for catalytic activity. Here, we found that the residues H275 and E333 in the C-cupin domain also had a significant influence on enzyme activity. Overall, both the C-cupin domain and N-cupin domain have a certain coordination effect on the catalytic process of OxDC.

To further verify the important role of the two residues (E162 and E333) interacting with Mn^2+^ in the two domains, respectively, we evaluated the metabolism of oxalate for WT OxDC and these two mutants at the cellular level. When OxDC was transfected into cells, exogenous OxDC could metabolize oxalic acid after being expressed by cells, reducing the toxic side effects of oxalic acid on HEK293 cells. We created a cytotoxic oxalate cellular environment, the successful transcription of exogenous genes was detected by the experimental methods of qRT-PCR and fluorescence microscope photographing (**Figure 5E**), and then found that the mRNA levels of apoptosis factor (Caspase3) and cellular inflammatory factors (IL-1β, TNFα) in HEK293^E162D^ and HEK293^E333D^ cells were higher than those in HEK293^WT OxDC^ cells (**Figure 5F**). This finding indicates that mutation of E162D and E333D could impair the oxalate-metabolization activity of OxDC.

### Discovered and verified the special effect of Zn^2+^ on OxDC

During the catalytic process, oxalate and dioxides bind Mn^2+^ to form a formyl radical anion intermediate to facilitate OxDC to metabolize oxalate to formate and CO_2_, thereby completing the decarboxylation reaction (40). Since OxDC is a metal-dependent enzyme, we explored the effects of common divalent metal ions on protein activity. The activity of OxDC in supplied with Ca^2+^, Mg^2+^, and Sr^2+^ were almost as same as the control group, indicating no effect on the activity of OxDC.

While the addition of Zn^2+^ and Mn^2+^ to OxDC retain the protein activity equally (**Figure 6A**). Given the good effect of Zn^2+^ on healthy intestinal flora, we emphatically analyzed Zn^2+^ for OxDC. The circular dichroism (CD) result revealed that Zn^2+^ generated similar effect on the secondary structure of the protein as Mn^2+^ did. Among them, α-helices decreased, while the proportion of β-sheets and β-turns was increased (**Figure 6B**), indicated that OxDC has the binding force to metal ions Mn^2+^ and Zn^2+^. Besides, Zn^2+^ improved the stability of the protein significantly (**Figure 6C**). ITC assay result showed that the binding capacity of Zn^2+^ to the EDTA treated OxDC was 130 times that of the untreated group, indicating that Zn^2+^ was likely to improve the thermostability and activity through competing with ‘Mn^2+^ to bind the active pocket of OxDC (**Figures 6D-E**).

**Figure 6.**
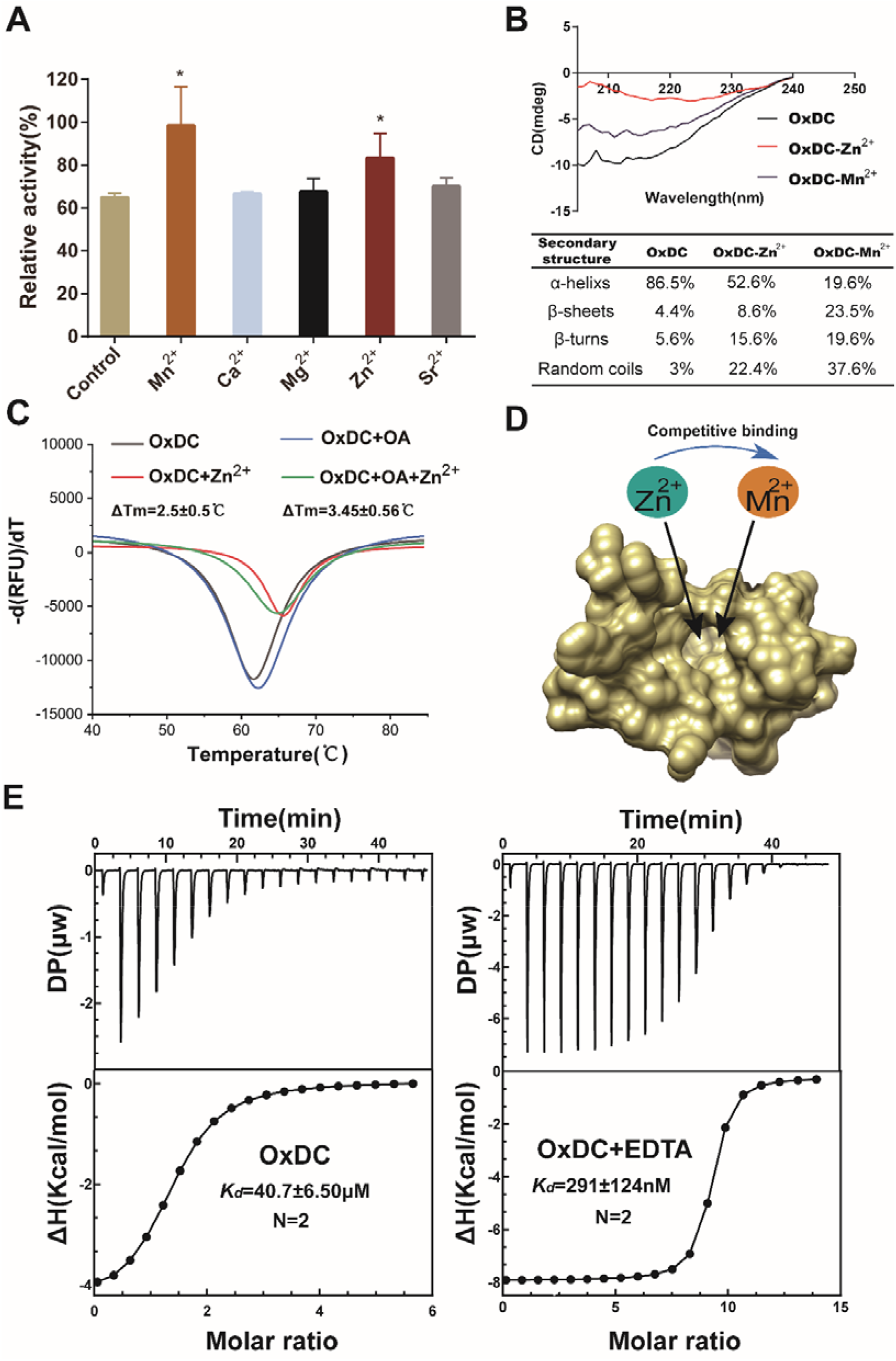
Discovered and verified the special effect of Zn^2+^ on OxDC. (A) Effects of common divalent metal ions on OxDC enzyme activity. The highest activity was set as 100% as a reference, the control group is the group without metal ion treatment and assays were carried out using a methylene blue-potassium dichromate system. (B) The effect of Zn^2+^ and Mn^2+^ on the secondary structure of OxDC tested by CD. (C) The influence of Zn^2+^ on OxDC’s thermal stability. (D) Schematic diagram of the binding pocket of Zn^2+^ competitively binding to Mn^2+^ (E) The binding ability of OxDC with Zn^2+^ under different treatment conditions was determined by ITC experiments. For processing of data from the protein activity experiment, * indicates *P*<0.05.

### The effect of Zn^2+^ on the digestive stability of OxDC *in vitro*

Previous studies have shown that the metal ion was important of OxDC activity, the supplementation of Zn^2+^ can increase OxDC’s activity and Mn^2+^ evenly distributed in the two domains of OxDC, which are crucial for metabolizing the oxalate. The result of chromatogram from size-exclusion chromatography analysis showed that OxDC exists in two major forms which are dimer and hexamer the functional aggregation state and 1mM EDTA reduce the hexamer of OxDC, because EDTA can attract metal ions likes Zn^2+^ and Mn^2+^ and induce the stability and activity of OxDC. When supplementing superfluous Zn^2+^ and Mn^2+^, the hexamer of OxDC can recover (**Figure 7A**). The same as OxDC protein thermal shift assay, add Zn^2+^ and Mn^2+^ can resist the reduce of Δ*T*_m_ from EDTA (**Figure 7B**). In addition, following previous experiment we detected the digestive stability of OxDC with trypsin *in vitro*. As displayed in **Figure 7C**, the signal of the OxDC band (∼44 kDa) decreased slowly as digestion time increased, indicating that the degradation of OxDC by trypsin was slight after 10 min and serious more than 30min. When the metal ions like Zn^2+^ and Mn^2+^ was superfluous, the major band of OxDC was recovered. In summary, metal ions were important for OxDC especially Mn^2+^ mentioned earlier. Our study found that Mn^2+^ can maintain the activity and structures stability. The activity of OxDC will be destroy, when Mn^2+^ attracted by EDTA. Adding superfluous Zn^2+^ can rebuild its activity and structures stability which improve the metabolism of Oxalic acid (**Figure 7D**).

**Figure 7.**
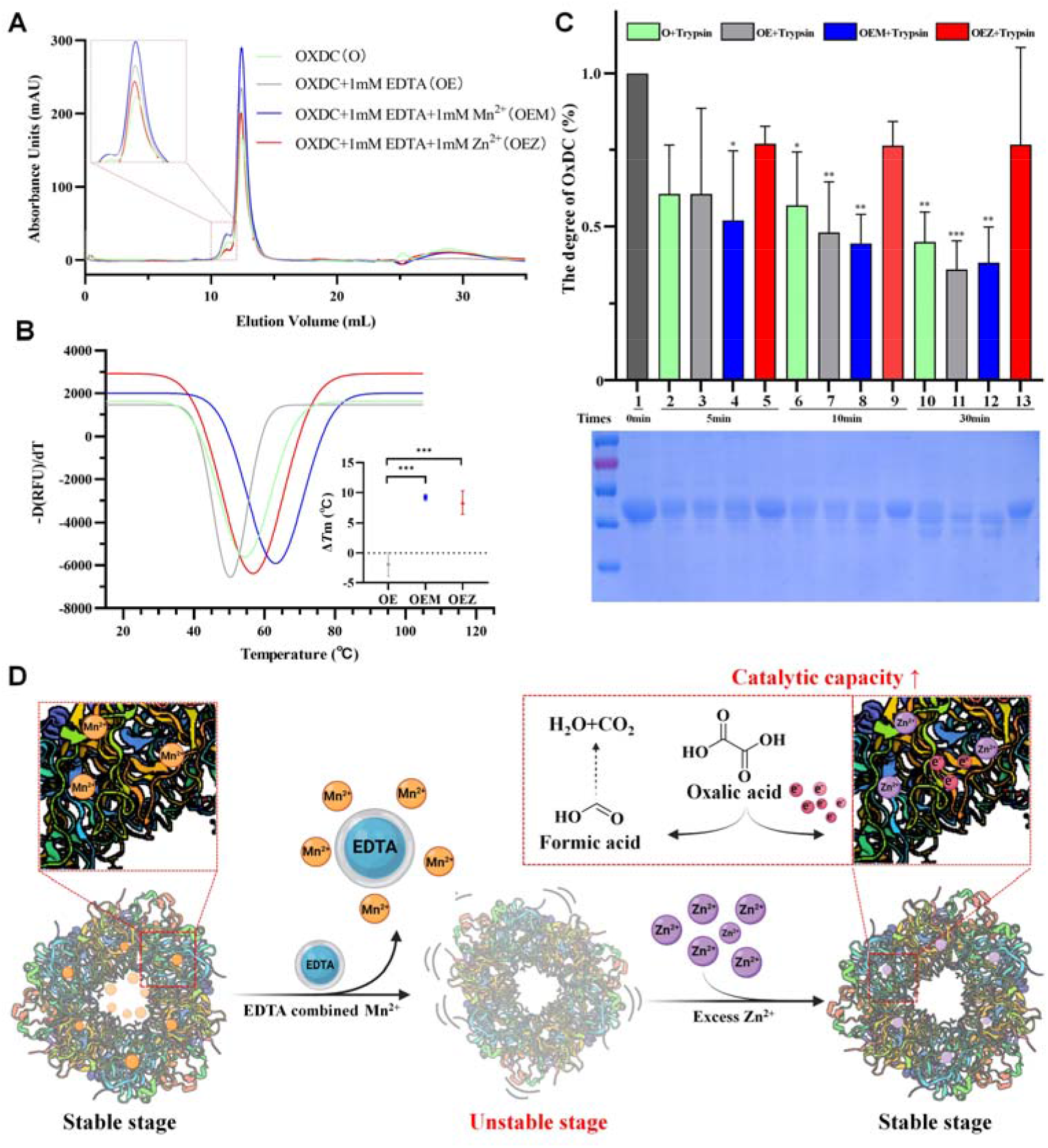
The effect of metal ion on the digestive stability of OxDC. (**A**) Chromatogram from size-exclusion chromatography analysis. (**B**) OxDC protein thermal shift assay. (**C**) *In vitro* digestive stability of OxDC with trypsin. (**D**) The sketch map which the important of metal ions Zn^2+^ and Mn^2+^ for OxDC.

## Discussion

In recent years, the relationship between oxalate-degrading bacteria and CaOx stones becomes research hot spots. In this study, we designed clinical trials and analyzed many clinical samples to further confirm that the *Lactobacillus* is closely related to the occurrence of CaOx stone and first found that the zine gluconate can increase the abundance of oxalate metabolizing bacteria in the intestine of CaOx stone patients. This finding shows that a reduction in the number of *Lactobacillus* may lead to a decline in the OxDC content, thereby reducing the metabolism of oxalate by the intestinal flora and increasing the incidence of CaOx stones, and supplementing zine gluconate has a positive effect on oxalate metabolizing bacteria.

Interestingly, we found that when conserved residues that interact with Mn^2+^ in the C-terminal cupin domain were mutated, the activity of the enzyme was also significantly decreased. This indicates that both metal sites are important and show synergy in the metabolism of oxalate. In addition, weak metabolic activity was still observed after key residues were mutated. This is related to the fact that OxDC can be converted into an oxalate oxidase under certain conditions. Because OxDC is likely to have evolved from oxalate oxidase by gene replication and exhibits strong sequence conservation with oxalate oxidase (41), however, the products of OxDC and oxalate oxidase are different, for example, oxalate oxidase can metabolize oxalate into hydrogen peroxide and carbon dioxide (42).

During the experiment, we found that Zn^2+^ has a protective effect on *Lactobacillus* under an antibiotic environment. According to reports, Zn^2+^ can be enriched by *Lactobacillus* by binding to peptidoglycan in the bacterial cell wall (21). Simultaneously, antibiotics mainly destroy the cell wall structure of bacteria, causing the cells to burst and die. We preliminarily believe that when Zn^2+^ is present around the *Lactobacillus*, Zn^2+^ will combine with the hydroxyl group, carboxyl group, and the other anions in the peptidoglycan structure of the bacterial cell wall to increase the density of the cell wall structure. Thereby *Lactobacillus* is immune to the interference of antibiotics, which might contribute to the protective effect of Zn^2+^ to *Lactobacillus*. OxDC is an Mn^2+^-dependent enzyme, and Mn^2+^ acts as a ligand for the redox cycle in the process of oxalate metabolism. Zn^2+^ and Mn^2+^ are two common divalent ions. We firstly found that Zn^2+^ can competitively bind to the Mn^2+^ binding pocket in OxDC, when all the metal ions contained in the protein itself were chelated by EDTA, the binding capacity of OxDC to Zn^2+^ was increased by about 100 times. This shows that Zn^2+^ can not only occupy the 12 binding sites of Mn^2+^ but may also occupy the binding sites of other metal ions, and it can increase the protein stability and change the secondary structure to enhance OxDC’s ability to bind and metabolize OA. At present, the precise relationship between Zn^2+^ and kidney stones is unknown. The earliest studies showed that the amount of urinary Zn^2+^ excreted from patients with stones mostly differs from that of healthy people (43), but this study was greatly affected by sex-related difference and individual difference. In a recent literature report, Mg^2+^ improved the survival rate of *B. subtilis* by regulating antibiotics that target the ribosome. We speculate that Zn^2+^ has a similar function. Therefore, we designed clinical trials and found that zine gluconate can indeed increase the allowance of *Lactobacillus* in the intestine of CaOx stone patients. However, the overall effect of zine gluconate on *Firmicutes* has little change. It is possible that zine gluconate not only increases the content of oxalate metabolizing bacteria, but also inhibits the abundance of some intestinal flora in *Firmicutes*, which needs to be further researched. In this paper, we explored the specific effect of Zn^2+^ on kidney stones from the perspective of *Lactobacillus* in the intestinal flora and OxDC originating from *Lactobacillus* for the first time, and our results provide a new strategy and direction for the prevention in stones via metal ions.

In recent years, among the four oxalate degrading bacteria, researchers have focused more on the relationship between CaOx stones and *Oxalobacter*. However, *Oxalobacter* is very sensitive to antibiotics and is present in the intestines of only 60%-80% of adults. Bariatric surgery, bowel disease, and cystic fibrosis all inhibit the recolonization of *Oxalobacter* in the intestine (44). In addition to its stability and high safety, *Lactobacillus* also has the strength of the intestinal barrier, body’s immune response, and play an important role in maintaining the balance of intestinal flora. These characteristics of *Lactobacillus* provide a theoretical basis for the development of related preparations to treat CaOx stones in the future.

Here, the structural basis and metabolic mechanism by which oxalate decarboxylase metabolizes oxalate were elucidated, and Zn^2+^ was illustrated to have therapeutic effects on CaOx stones by improving the tolerance of *Lactobacillus* to antibiotics. According to our research, proper Zn^2+^ levels in the diet, the consumption of more probiotic food and avoidance of the antibiotic overuse might be desirable measures for the prevention and treatment of kidney stones.

## Methods

### Materials and reagents

The OxDC gene derived from *L. farciminis* (GenBank accession number: CP012177.1) was constructed by GenScript company (Nanjing, Jiangsu, China) and cloned into the pET-20b (-) vector. *E. coli* DH5α and BL21 (DE3) were the host strains for plasmid amplification and protein expression, respectively. The *Nde*I and *Xho*I restriction enzymes were purchased from Thermo scientific. Ni-NTA column, HisTrap HP 1mL and Superdex 200 increase 10/300GL resin were all purchased from GE Healthcare. SYPRO Orange protein gel staining 5000x and PrimeScriptTM RT reagent Kit were purchased from Thermo Fisher Scientific. *L. casei* was a gift from Macau University of Science and Technology. oxalic acid, sodium oxalate, NH_4_Cl, and EG were purchased from Sigma-Aldrich. Chloramphenicol, clindamycin, vancomycin, ampicillin, kanamycin, penicillin, gentamicin was supplied by Ruishu Bio. Zinc Gluconate was purchased from the hospital. Other chemicals used were biochemical research grade. *L. casei* was selected for all *Lactobacillus* assays in this study.

### Patients and samples

This research was approved by the Human Ethics Committee of Guangdong Provincial Hospital of Chinese Medicine (Guangzhou, Guangdong, China). An informed consent form was signed by all patients in the study (ZF2020-059-01). The clinical data and samples were harvested from the Second Affiliated Hospital of the Guangzhou University of Chinese Medicine (Guangzhou, Guangdong, China) from July 2018 to January 2022. The inclusion criteria for patients in figure 1 were as follows: 1) diagnosis of CaOx kidney stones, 2) age of 20-60 years, 3) glomerular filtration rate below 60 ml/min/1.73 m^2^, 4) no uptake of drugs or food to regulate the intestine within the past one month, and 5) no significant changes in diet. The exclusion criteria were as follows: 1) stones with the tumor or bladder stones, 2) hypotension, hypernatremia, or a history of congestive heart failure, 3) pregnancy, 4) hyperthyroidism, adrenal diseases, pituitary tumors, or other diseases that affect calcium regulation, 5) present history of intestinal disease or history of major gastrointestinal surgery, 6) serious heart, brain, liver, kidney or other systemic diseases or mental illness, or 7) severe urinary tract infection. In Figure 1, we divided the subjects into antibiotic treatment group, the antibiotic combines with Zinc Gluconate group, the patients with CaOx Kidney stones group and the patients with No-kidney stones group. Each group contained ten subjects. Using 16SRNA technology to measure the stool of patients, and the subjects had good compliance. The CaOx composition of the stones was confirmed by infrared spectrum analysis. Feces were provided as a specimen of all participants.

### The influence of external factors on the growth of *L. casei*

MRS medium was subjected to high-temperature steam sterilization and used as the culture medium for *L. casei*, a strain of *Lactobacillus*. Through drug sensitivity assays, we examined the effects of seven antibiotics (chloramphenicol, kanamycin, ampicillin, vancomycin, clindamycin, penicillin, and gentamycin) on *L. casei* growth. *L. casei* was cultured in MRS medium containing antibiotic at a final concentration of 0.1 μg/ml. To explore the effect of Zn^2+^ on the oxalic acid tolerance of *L. casei*, oxalic acid was added at a three-concentration gradient (1, 5, and 10 mM) and 2 mM Zn^2+^ to the MRS medium used to culture *L. casei*. To test the effects of Zn^2+^ supplementation on the antibiotic tolerance of *L. casei*, 0.8 μg/ml antibiotic and Zn^2+^ at 2 mM and 10 mM were used. The absorbance was measured at OD_600_ with a microplate reader at fixed time points. This method was based on previously reported literature (45).

### Animal experiments

Six-week-old male Sprague-Dawley (SD) rats were purchased from Southern Medical University, and the animal protocol was approved by the Animal Care Committee (SYXK2019-0144). The SD rats were housed at 22 ± 2 °C in standard cages in a specific pathogen-free facility under a 12-h and 12-h light-dark cycle. Water and chow were available ad libitum. After the weights of the SD rats were measured, they were randomly divided into 5 groups containing 5 rats each. The rats in control group received normal rat chow and drinking water. Rats in groups - “Stone”, “Amp+Stone”, and “Amp+Zn^2+^+Stone” were received food containing 0.4% NH_4_Cl and 1% EG from day 14 to day 29 to form CaOx kidney stones. Rats in group- “Amp+Zn^2+^+Stone” was received supplemental Zn^2+^ for 29 d, while rats in group-“Amp+Stone” and “Amp+Zn^2+^+Stone” were received antibiotics for 29 d. Amp and ZnSO_4_ were dissolved in ultrapure water and administered by gavage to the rats every day at 9:00 and 16:00 at doses of 20 mg/kg and 10 mg/kg (body weight). Kidney, urine, and blood were collected on days 3, 5, 7, 9, 11, 13, and 15 during the experiment. The body weight of the animals was monitored, and a urine sample (24 h) was used to detect pH. The kidney was subjected to H&E staining to observe the growth of the stone, and the collected serum was used to measure the Zn^2+^ content. All the animal experiments involved in this study were approved by the Institutional Animal Care and Use Committee of the Guangzhou University of Chinese Medicine.

We choose the EG to build a CaOx kidney stones model, and this method consists of the administration of 0.4% NH_4_Cl and 1% EG, the precursor of oxalic acid, which increases the concentration of uric acid, while NH_4_Cl acidifies the urine and accelerate the growth of stones (46). NH_4_Cl and EG dissolved in ultrapure water to final concentrations of 0.4% and 1%, respectively, were administered in the drinking water, which was freely available to the rats for 15 d. The rats were dissected and assessed for kidney stones growth on the 14^th^, 17^th^, 19^th^, 21^st^, 23^rd^, 26^th^, and 29^th^.

### Hematoxylin-eosin staining

After the kidney tissues were washed with 0.9% normal saline, they were first fixed with 4% paraformaldehyde for 48 h and then dehydrated, waxed, and paraffin sectioned to a thickness of 4 μm. The sections were further dewaxed with xylene and eluted with a gradient of ethanol. Finally, the sections were dehydrated, permeabilized and sealed by dying with Yin Hong staining for 2 min, and finally dehydrated with ethanol, then permeabilized and sealed, completing the HE stains process. Prepared tissue sections were observed under an optical microscope at a 200× magnification.

### Determination of urea nitrogen in serum and urine

Centrifuge the collected rat blood samples at 4°C, 3000 rpm for 10 min, and collect the supernatant. Dilute the collected serum samples by 5 times, and then use the urea nitrogen detection kit for content determination. Urea is hydrolyzed under the action of adenase to produce NH_4_^+^ and CO_2_. NH_4_^+^ generates blue substance with phenol developer in alkaline environment. The amount of blue substance produced is proportional to the content of urea nitrogen, which is used as a calculation. Similarly, the collected rat urine was diluted 5 times and tested in the same way. Finally, calculate according to the formula on the manual.

### Collection of stool samples, 16S RNA sequencing, and data analysis

To detect the changes in the intestinal flora of the four groups of volunteers recruited, we collected volunteer feces on a regular basis. Professional collection was performed by nurses with the permission of volunteers. Finally, the collected samples were sent to Majorbio for determination.

### Construction of vectors

We constructed a truncated protein consisting of the main body of *L. farciminis* OxDC (aa 22-373, WT) using recombinant cloning technology and obtained five mutant plasmids (H95Q, E162D, H275Q, E333D, and Y340N) using *QuikChange* technology. The truncated OxDC gene was subcloned into the pET-21b (-) vector with a C-terminal 6×His tag, and the WT and mutant genes were subcloned into the pET-20b (-) vector. PCR-amplified WT, E162D and E333D fragments were cloned into the pEGFP vector at the *Nhe*I and *BamH*I restriction sites and expressed in the eukaryotic expression vector pEGFP. All recombinant plasmids were confirmed by PCR and DNA sequencing.

### Protein expression and purification

The proteins were overexpressed in the *E. coli* BL21 (DE3) and Rosetta (DE3) strains. Bacterial cultures were grown in LB medium at 37 °C. When the OD_600_ was 0.6-0.8, cells were induced with isopropyl β-D-1-thiogalactopyranoside to a working concentration of 0.2 mM. After incubation at 25 °C overnight, *E. coli* cells were centrifuged for 20 min at 4,000 rpm, and cell lysates were obtained with a cryogenic grinder and then centrifuged at 14,000 rpm for 1 h at 4 °C to obtain the supernatant. The resulting supernatant was roughly purified using Ni-NTA agarose resin (GE Healthcare, Sweden). Further purification method was carried out with a 1 mL HisTrap HP column (GE Healthcare, Sweden) to obtain the pure protein. Since crystals of the full-length protein do not diffract, the truncated protein corresponding to the body of OxDC (aa 22-373) was prepared for the crystallographic experiment. All proteins were purified in a similar manner.

### Size-exclusion chromatography analysis

Using an AKTA pure 25 M1 (GE Healthcare, Sweden) purification system with Superdex 200 increase 10/300 GL resin column, the state of the protein in solution was examined. The target protein was centrifuged at 14,000 rpm for 10 min at 4 °C before loxalic acidding, and the eluted solution contained 20 mM Tris-HCl (pH 8.0), 150 mM NaCl, and 1 mM DTT. Three protein standards (440 kDa, 158 kDa, 75 kDa) were dissolved together in the same solution as target protein and separated by gel filtration chromatography.

### Crystallization, data collection, and structure determination

Purified OxDC was concentrated to approximately 3.5 mg/ml. The initial crystal was obtained using a commercial screening kit (Index) with the 96 well sitting-drop vapor diffusion method. In the optimization process, 2 mM Mn^2+^ and 10 mM oxalate were incubated with the protein in an acidic environment for 1 h at 4 °C. The protein and the reservoir solution were mixed in a 2:1 volumetric ratio. The best crystals were produced in a reservoir solution containing 18% PEG3350, 0.2 M NH_4_Cl, 0.2 M sodium dihydrogen phosphate, and 0.2 M Bis-Tris (pH 5.5). The selected crystals were soxalic acidked for 1-3 min in a protective solution containing all the components of the reservoir solution supplemented with 20% glycerol (v/v), and crystals were then immediately flash frozen in liquid nitrogen. Data collection was performed at beamline 17U (BL17U1) of the Shanghai Synchrotron Radiation Facility (SSRF, Shanghai, People’s Republic of China), and data processing was carried out with the program HKL2000. Molecular replacement was performed with CCP4 software using the known structure of *B. subtilis* OxDC (PDB ID: 1UW8) as a search model. By further build model manually with Coot software (47), the structure of *L. farciminis* OxDC at a resolution of 2.79 Å was finally obtained.

### Cell culture and treatment

HEK293 cells were purchased from China Center for Type Culture Collection (CCTCC) and cultured in Dulbecco’s modified Eagle’s medium (DMEM) supplemented with 10% fetal bovine serum, 100 U/ml penicillin and 0.1 mg/ml streptomycin. The incubation conditions were 37 °C, 95% humidity and 5% CO_2_. Transfection of cells at a specific stage of cell culture was performed according to the instructions of a transfection kit (Hanbio, China). After 24 h, transfectants with a fluorescence signal appeared. The overexpression of foreign genes was further confirmed by qRT-PCR.

Cell viability was evaluated by the MTT method. HEK293 cells expressing recombinant pEGFP, pEGFP^WT^, pEGFP^E162D^ and pEGFP^E333D^ were treated with 0 μmol, 50 μmol, 250 μmol, 500 μmol, 750 μmol, or 1000 μmol of oxalate. After the cells were further cultured, MTT was added to the cell medium at a final concentration of 0.5 mg/ml and incubated for 4 h. The cell culture plate was removed from the incubator, 200 μl of DMSO was added to dissolve the crystal violet, and the absorbance was then immediately recorded at OD_570_ after incubation on a shaker in the dark for 5 min. The optical density of each well was subtracted from blank control well. After analysis of the results, 750 μM oxalate was selected to generate a toxic oxalate environment.

### *In vitro* enzyme activity assay

The enzymatic activities of the WT and mutant OxDC proteins were examined by methylene blue-potassium dichromate assay. Pure proteins containing 5% glycerol were mixed with oxalic acid at a molar ratio of 1:100, and all enzymatic solutions were incubated in a reaction system of 100 mM NaCl and 40 mM Bis-Tris (pH 5.5) for 2 h at room temperature. The reaction was terminated by the addition of 600 μl of 1 M H_2_SO_4_. Then, the remaining concentration of oxalic acid in the enzyme reaction system was determined immediately by methylene blue-potassium dichromate redox catalysis. Finally, the absorbance of the reaction solution at OD_660_ was measured, and a standard linear relationship was established between the oxalic acid concentration and the absorption value. All assay results were repeated five times or more.

### Protein thermal shift

SYPRO Orange protein gel staining 5000× (Thermo scientific, USA) was diluted to 200x with 40 mM Bis-Tris (pH 5.5), 100 mM NaCl, and 1 mM DTT. All protein samples were concentrated to about 4 mg/ml. The assay contains two groups, one group was used to investigate the effect of adding Zn^2+^ on OxDC, and the other group was used to explore the effect of adding Zn^2+^ on OxDC after incubating with 2 mM oxalic acid. 2 mM Zn^2+^ was incubated with the protein for 1 h, and then centrifuged it at 14000 rpm for 10 min for detection. The total 20 μl reaction system was mixed with 5 μl SYPRO Orange protein gel staining 200× and 20 μg protein. The sample was gradually heated from 25 to 90 °C at a gradient of 0.2 °C /min and analyzed by qRT-PCR instrument. All protein thermal shift figures were produced by Origin, version 19.0.

### Circular dichroism (CD) assay

The purified protein was concentrated to 1 mg/ml in 20 mM Tris 8.0, 100 mM NaCl. Then, 1 mM Zn^2+^ and Mn^2+^ were added to 200 μg of protein incubated on ice for 1 h. Set the wavelength to 180-260 nm for scanning, observe the chromatographic peaks and use CDpro software for data analysis and processing. Finally, the changes in the secondary structure of the protein are judged by comparing the ratio changes of the indicators (*α*-helixs, *β*-sheets, *β*-turns and random coils).

### Isothermal titration calorimetry (ITC) assay

The ITC experiment was carried out using Auto-ITC (Malvern, United Kingdom) instrument at a constant temperature of 25 °C. During setting up the program, the volume of the first drop of small molecules is 0.4 μl, and the volume of the remaining 18 drops is 2 μl per drop. After the experiment, use MicroCal ORIGIN software to process and analyze the data. When investigating the binding ability of OxDC and oxalic acid, 10 mM oxalic acid (60 μl) was used to titrate 340 μM (200 μl) OxDC. During investigating the binding mode of Zn^2+^ and OxDC, firstly, the purified protein was dialyzed into a buffer containing 100 mM NaCl, 20 mM Bis-Tris (pH 6.0) and 20 mM EDTA. After 2 h of dialysis, multiple dialysis methods were used to remove the ions. The protein was dialyzed into 100 mM NaCl, 20 mM Bis-Tris 6.0 buffer to facilitate the removal of EDTA. The control protein does not need to be pre-treated with EDTA.

### Digestive stability of OxDC *in vitro*

To determine if the Zn^2+^ and Mn^2+^ affect the stability of OxDC enzyme, we performed susceptibility to protease assays. The OxDC protein was adjusted at 1.0 mg/mL in Tris-HCl buffer (20 mM Tris-HCl, pH 7.0), then incubated with 1mM EDTA and 1mM Zn^2+^ or Mn^2+^, at last incubated with 1mM Trypsin for 1, 5, 15 min at 25°C. The quality ratio of OxDC to Trypsin was 500. Finally, digestive stability of OxDC was verified by 15% sodium dodecyl sulfate-polyacrylamide gel electrophoresis (SDS-PAGE) and Colloidal Brilliant Coomassie Blue R-250 was used to stain the gel.

### Statistical analyses

All the grouped data were assessed by GraphPad Prism software 7.0 (GraphPad Software, USA). Data were statistically calculated using t-test (two groups) or one-way ANOVA. Independent experiments were performed at least five times with similar results were considered significant if *P* values <0.05.

## Acknowledgments

This work was supported in part by the Guangdong Key Laboratory for Translational Cancer Research of Chinese Medicine (No. 2018B030322011), Guangdong Province Universities and Colleges Pearl River Scholar Funded Scheme (No. Caiyan Wang, 2019), Natural Science Foundation of Guangdong Province (No. 2017A030310501), and Scientific Research Project Funded by Traditional Chinese Medicine Bureau of Guangdong Province (No. 20221170).

## Author contributions

Caiyan Wang and Zhongqiu Liu conceived the project and designed the experiments; Fang Wu, Yuanyuan Cheng, Peisen Ye, Xuehua Liu and Lin Zhang carried out experiments and analyzed the data in this article; Yuanyuan Cheng, Fang Wu, Jianfu Zhou and Rongwu Lin wrote the paper; Songtao Xiang and Caiyan Wang advised and revised the manuscript. All authors approved the final version of the manuscript, and all the authors declared no competing interests.

## Declaration of Competing Interest

The authors have declared that no competing interest exists.

